# Detection of EGFR and its Activity State in Plasma CD63-EVs from Glioblastoma Patients: Rapid Profiling using an Anion Exchange Membrane Sensor

**DOI:** 10.1101/2023.10.16.562628

**Authors:** Nalin H. Maniya, Sonu Kumar, Jeffrey L. Franklin, James N. Higginbotham, Andrew M Scott, Hui K Gan, Robert J. Coffey, Satyajyoti Senapati, Hsueh-Chia Chang

## Abstract

We present a novel quantitative immunoassay for CD63 EVs (extracellular vesicles) and a constituent surface cargo, EGFR and its activity state, that provides a sensitive, selective, fluorophore-free and rapid alternative to current EV-based diagnostic methods. Our sensing design utilizes a charge-gating strategy, with a hydrophilic anion exchange membrane and a charged silica nanoparticle reporter. With sensitivity and robustness enhancement by the ion-depletion action of the membrane, this hydrophilic design with charged reporters minimizes interference from dispersed proteins and fluorophore degradation, thus enabling direct plasma analysis. With a limit of detection of 30 EVs/μL and a high relative sensitivity of 0.01% for targeted proteomic subfractions, our assay enables accurate quantification of the EV marker, CD63, with colocalized EGFR by an operator/sample insensitive universal normalized calibration. Glioblastoma necessitates improved non-invasive diagnostic approaches for early detection and monitoring. Notably, we target both total and “active” EGFR on EVs; with a monoclonal antibody mAb806 that recognizes a normally hidden epitope on overexpressed or mutant variant III EGFR. This approach offers direct glioblastoma detection from untreated human patient samples. Analysis of glioblastoma clinical samples yielded an area-under-the-curve (AUC) value of 0.99 and low p-value of 0.000033, significantly surpassing the performance of existing assays and markers.

## 1. Introduction

Liquid biopsy is an emerging non-invasive approach for detecting circulating cancer biomarkers in various body fluids^1^. Extracellular vesicles (EVs), including exosomes and microvesicles, have gained interest as a target for cancer detection^2^. EVs play a role in paracrine cell signalling as nanocarriers for the exchanged RNAs and proteins^3^. They possess the ability to transport highly charged and hydrophilic molecules across the hydrophobic bilayers of cell membranes, and they provide protection to miRNA and mRNA from degrading agents^1^. Furthermore, EVs and their cargo are excreted abundantly and exhibit high stability in various body fluids, such as blood, urine, and saliva^4^. This stability and abundance have motivated the development of diagnostic assays based on EV biomolecular contents.

Here, we focus on one such aspect – the tumour-specific epitope on the CR1 domain of the Epidermal Growth Factor Receptor (EGFR) present on the surface EGFR-amplified cells^5^ and their secreted EVs^6^. Under normal conditions in healthy cells, this epitope is transient and mostly hidden due to a disulfide-bonded loop between amino acids 287 and 302 on EGFR, which creates steric hindrance with the CR1 region^7^. However, in EGFR-amplified cancers, the disruption of this bond in multiple EGFR copies reduces this steric hindrance, making the epitope accessible in the untethered EGFR^7^. A similar exposure of the epitope occurs in the mutant variant III EGFR^8^, where the deletion of amino acids 6-273 exposes the same epitope due to the significant deletion of L1 and CR1 domain. Therefore, an antibody (mAb 806) specifically targeting this sterically hidden epitope would be ideal for detecting cancer EVs secreted by EGFR-amplified tumour cells^9^, as the same active version of EGFR (aEGFR) is also shared on the tumour-secreted EVs^6^. Other EGFR antibodies such as cetuximab and panitumumab are distinct from mAb 806 as they recognize total wtEGFR (tEGFR)^10^.

However, a key challenge associated with detecting aEGFR using mAb 806 is the relatively high dissociation constant^7^ *K*_*D*_∼30 *nM*, while the concentration of aEGFR in plasma is much lower with overall tEGFR concentration <1-10 pM^11^. Most of the current approaches for examining proteins on EVs, such as Enzyme Linked Immunosorbent Assay (ELISA) and immunoblotting involve lysis of EVs. These methods typically have a limit of detection that is 10-100 times lower than *K*_*D*_^12, 13^ and hence insufficient for robust detection of dispersed aEGFR from lysed EVs. Furthermore, these approaches often require laborious ultracentrifugation and enrichment steps to visualize a band, which becomes even more difficult when working with human plasma due to the presence of high contaminant from non-EV species. Efforts to isolate EV also lead to variation due to significant EV loss (∼90%) when high-force ultrafiltration or ultracentrifugation is used to isolate them from other proteins and lipoproteins^14^ leading to a yield bias. Similarly, fluorescent labelling of aEGFR on EVs for flow cytometry or nanoparticle tracking analysis requires EV isolation from plasma to remove unincorporated fluorophores and overcome high labeling interference from dispersed proteins and reactive oxidative species^15^. Autofluorescence from abundant dispersed proteins, such as albumin, is also an issue for any optical assay of raw plasma that involves fluorescent labelling^16, 17^.

An alternative approach to quantify aEGFR is to capture EVs with a tetraspanin EV marker (CD63) with high capture affinity and report it with anti-aEGFR at very high reporter concentrations ≫ *K*_*D*_, leading to a sandwich scheme with irreversible antibody association with the *colocalized protein* (aEGFR). However, due to endogenous and handling-induced variations in total EV number, absolute quantification of aEGFR+ EV leads to poor p-statistics. Instead, capturing a fraction of CD-63 EVs with aEGFR colocalized in untreated plasma sample would reduce false positives and negatives due to EV number fluctuations. This normalized colocalization assay requires a large sensor dynamic range because the dynamic range determines the lowest colocalized fraction that can be accurately determined. When captured with tetraspanins and reported with a particular protein of interest, the signal is essentially below the limit of detection if the colocalized fraction is only 1% of the captured EVs in a sensor with a 2log_10_ dynamic range.

Therefore, immuno-sandwich characterization of aEGFR-positive EVs necessitates three key requirements. First and foremost, a high sensitivity detection method is essential to accurately detect the low concentration of aEGFR-EVs or any other EV subfractions in plasma. Secondly, the sensor employed should possess a large dynamic range to enable the reliable determination of colocalized fractions even at low levels. Finally, it is crucial to develop an interference-free approach that eliminates the need for laborious EV isolation procedures. By fulfilling these requirements, the detection and characterization of aEGFR-positive EVs in untreated plasma can facilitate advancements in liquid biopsy-based cancer diagnostics. Many of the recently proposed immuno-sandwich colocalization assays fail to meet these requirements. Interferometry-based colocalization assays^18^ typically only exhibits one-log dynamic range and hence cannot accurately estimate a colocalization fraction below 10 percent. Electrochemical assays often require EV isolation due to interference from abundant redox agents in plasma^19-21^.

We report here the first normalized non-optical colocalization assay for untreated plasma EVs, with sufficient sensitivity (30 EVs/μ*L* of sample) and selectivity to quantify the fraction of CD63 EVs with a disease marker in untreated plasma. We use a charge sensing strategy with a highly charged silica nanoparticle reporter to minimize signals from non-target molecules and EVs, which are typically weakly charged. The dimension of the charged reporter (50 nm) is selected such that only a single reporter can bind with one target EV, due to steric and electrostatic repulsion, such that the charge signal is identical for EVs of different size. The sensor utilizes the long-range (∼1 mm) ion depletion action of an ion exchange membrane^22-24^ to amplify signal transduction by minimizing Debye screening of the reporter (EV size varies by 100 nm and the plasma Debye screening length is less than 1 nm). With depletion, the field of charged reporter can reach the membrane surface and gate the ionic current, independent of its distance to the membrane due to EV size variation. Since the reporter signal is only registered after ion-depletion, it can be accurately quantified by a sensitive transition voltage to the “over-limiting” regime due to an electroconvective instability that occurs because of ion depletion^24, 25^.

The platform takes advantage of the strong binding affinity of antibodies towards CD63, which is further enhanced by avidity through a combination of high probe density and the presence of multiple copies of CD63 on the surface of EVs. This optimized capture strategy allows for high-affinity capture of a large number of EVs, maximizing the sensitivity of our assay. Unlike ELISA and immunoblotting, which require over 10^9^ reporters to detect a signal, our platform achieves significant sensitivity with only 10-1000 silica reporters. Additionally, the small dimension of our sensor (< 1 mm) offers the advantage of shorter incubation times for signal registration, especially at very low concentrations^24, 26^. Traditional surface assays with larger sensors often suffer from transport limitations and necessitate a larger number of reporters, resulting in prolonged incubation times before any signal can be detected. In contrast, our platform’s lower reporter requirement allows for faster signal registration, typically within a timeframe of approximately 30 minutes to 1 hour. This expedited process enables efficient signal detection even at low concentrations. We have also demonstrated that, after the diffusion time of 30 minutes for our small sensors, the total captured analyte does not change significantly, as diffusive depletion of the analyte is confined to a small micro-liter neighbourhood of the sensor^26^.

Once captured, the EVs are exposed to a high abundance of silica reporters that target either aEGFR, tEGFR (total EGFR) or CD63. This abundance of reporters ensures that all available binding sites on the captured EVs with our target protein are engaged, leading to rapid irreversible reporter binding within minutes. High reporter concentration overcomes low affinity or low concentration of target proteins on the EVs. Moreover, with the high probe density enabled by dense active sites on the membrane, the sensor can accommodate approximately 10^7^ silica reporters until saturation is reached^26^. This allows for a wide dynamic range of detection, spanning four to five logs. The signal increases proportionally with the logarithmic concentration of EVs, ranging from 100 to 10^7^ silica reporters. Therefore, our platform can accurately detect and quantify EV subfraction concentrations, even at levels as low as 0.01% experimentally. Finally, we also use the large drag force (several orders higher than on molecular analytes) on EVs and the reporter nanoparticle^27, 28^ to improve specificity with an optimized wash protocol. The engineered high sensitivity and selectivity of our sensor allow precise analysis of EV subpopulations within complex biological samples. By reliably detecting and quantifying EVs at low concentrations, our platform can detect rare subfractions of EVs, which can have significant implications in various fields including cancer research, biomarker discovery, and disease monitoring.

We validate this new AEM EV immunoassay by estimating the fraction of CD63 EVs with aEGFR in a small pilot cohort. We confirm that, due to irreversible association with the abundant reporters, a universal calibration curve exists that allows a very simple estimate of this fraction from the signals with two different reporters for two different colocalized proteins. We also validate the universality of this normalized colocalization assay with an HDL assay for colocalized ApoA1 and PON1. Finally, we analyze glioblastoma clinical samples and assess the total (tEGFR) and aEGFR fractions in CD63 EVs. With samples from both glioblastoma patients and healthy individuals, we also analyze the scatter of the data and arrive at a p-value of 0.000033 for aEGFR-CD63 EVs and 0.01 for tEGFR-CD63 EVs, each with an AUC of 0.99 and 0.755 respectively, both significantly better than any single-marker EV diagnostic assay with untreated plasma. The AEM sensor has been already verified for robust multiplexed quantification of RNA, DNA, proteins, and lipoproteins in our earlier work^26, 28-30^. However, it is particularly relevant to EV diagnostics due to the larger dimension and size variation of EVs. Additionally, we report here its fully automated prototype with near turn-key functionality.

## 2. Results and Discussion

### 2.1. Sensing Platform

The anion exchange membranes (AEM) display characteristic non-linear ion current-voltage (I-V) characteristics due to the ion-depletion action on one surface of the membrane which has been functionalized with the capture antibody^23, 25^. The overall architecture of our platform is shown in **Figure 1a**. Schematic I-V curves are shown in **Figure 1b**, with a linear Ohmic region at low voltage, followed by a high resistance limiting current regime beyond a critical voltage and then a third overlimiting current regime at an even higher transition voltage, with a differential resistance comparable to the Ohmic region. The automated prototype and algorithm is shown in **Figure 1c**. The ion-depletion action is highest at the overlimiting current transition voltage, and it is this transition voltage that is sensitive to the presence of charges on the membrane, and hence registers charge introduced by our capture antibody – EV – reporter antibody – silica nanoparticle complex. The ion-depletion action also ensures the measurements are always made at the same low ionic strength and neutral pH, independent of the sample ionic strength and pH. It hence introduces a universal deionized sensing buffer without extraction. This makes the sensor ideal for heterogeneous plasma or serum sample analysis as the sensor is insensitive to weakly or non-charged molecules like albumin. Even though these molecules non-specifically adsorb onto the sensor surface, the sensor does not register any signal as the sensor is only sensitive to charge. Of note, excessive fouling by uncharged non-targets will obviously still interfere with the sensitivity and so high probe density with large dynamic range is also required. The sensor measurements are performed at a constant current and the quantification signal is the voltage shift at that current.

**Figure 1.**
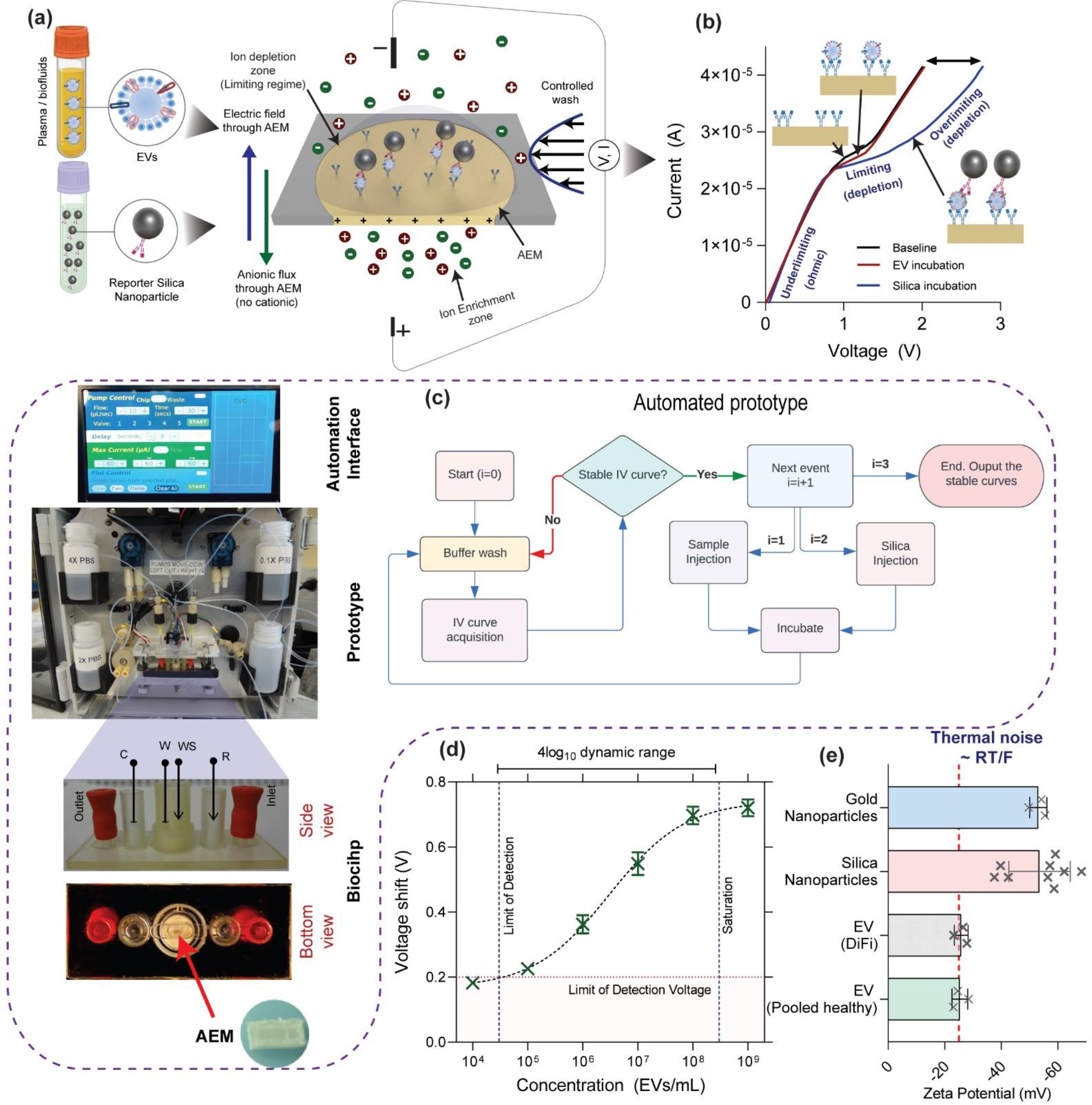
(a) General schematic and overall workflow of our platform. First, samples containing EVs are incubated, washed, incubated with silica reporters, and washed again. The platform consists of an anion exchange membrane that only allows counter ions to pass through and exhibits three distinct regimes in the current-voltage response. (b) Current-voltage response of an Anion Exchange Membrane: EV does not produce any shift in the overlimiting region, while silica reporters do produce a shift after forming a sandwich due to their highly negative charge. (c) Automation algorithm of our platform with an automation interface, prototype, and biochip showing the housing of AEM. (d) Voltage shift for anti-CD63 capture and anti-CD63 reporter (both monoclonal and of the same clone) that target a specific epitope of CD63 – binding only to species that contain at least two unique copies of CD63. (e) Highly negative zeta potential of silica compared to EVs from DiFi cells and pooled healthy plasma.

It is apparent from **Figure 1b** that when EVs bind to the anti-CD63 antibody on the sensor, a voltage shift is not observed. This is because EVs are weakly charged, with a Zeta potential of about -20 mV (**Figure 1e**) under low ionic concentrations (near DI conditions). However, as the silica nanoparticles functionalized with a reporter antibody are highly charged^31^ (Zeta potential ∼ -50 mV) (**Figure 1e**), resulting in a shift when it binds to the Evs on the sensor (**Figure 1d**) which can be compared against varying bulk concentration to establish a one-to-one correspondence between bulk EV concentration and voltage shift (discussed in detail later). We note that the Zeta potential scales as the logarithm of the charge density, but the total charge is the charge density multiplied by the surface area. Thus, we expect the reporter to introduce at least a few orders of magnitude more charge per EV. Additionally, the wash is done for 10-100 seconds with PBS which is the typical 1/*k*_*off*_ of non-specifically bound non-targets, while the shear force also removes non-specific large targets (such as non-target EVs and aggregates) thus using shear allowing us to significantly improve our selectivity.

### 2.2. Detection of Extracellular Vesicles

EV detection was performed in a sandwich scheme using a capture antibody attached to the AEM sensor (Figure S1, Supporting Information) and a reporter antibody attached to silica particles. We target the same specific epitope on CD63 using both capture and reporter antibodies due to the high abundance of tetraspanin CD63 marker on EVs secreted by most cell types^2, 32^. The concentration of reporter antibodies was maintained at 100 nM, to ensure it is higher than *K*_*D*_ of high-affinity antibodies. This means that a conservative *k*_*on*_∼10^5^*M*^−1^*s*^−1^, irreversible reporter association is reached within an incubation time of (*k*_*on*_*C*_*reporter*_)^−1^∼100*s*. Moreover, the attachment of antibodies to the AEM sensor was confirmed by using Alexa fluor 488 labeled antibodies. The bright green fluorescence observed from the sensor surface (Figure S2a, Supporting Information) indicates the successful attachment of the antibodies to the surface. Further, the labeled reporter antibody was used to confirm the attachment of antibodies to the silica particles (Figure S2b, Supporting Information). The presence of fluorescence only in the pelleted silica particles shows the efficient functionalization of the antibodies on the silica reporters.

To establish the one-to-one correspondence between the voltage shift observed in the overlimiting regime and the bulk EV concentration we use isolated EVs from DiFi media at increasing concentrations from 10^4^ -10^9^ EVs/mL (Figure S3, Supporting Information); in these studies, only one 50-nm reporter can associate with a single EV due to steric and electrostatic repulsion. First, the EV sample was incubated over the biochip for 20 minutes and then incubated with the reporter for 20 minutes. After each incubation step, there is a wash step. As seen from **Figure 1b**, the IV curves remain the same as the baseline curve with almost no voltage shift after EV binding. The binding of reporter particles caused the drastic voltage shift in the over-limiting region as expected from their zeta potentials. Here, the under-limiting and limiting regions remained constant which can be used as internal control for each sensor response. The blank is obtained by incubating PBS (with no EVs) followed by a silica reporter and is used to determine the limit of detection (LOD) voltage ∼ 0.2 V. The LOD of the AEM sensor was 30000 EVs/mL (quantified by NTA) which is more than 1000-fold lower than the conventional ELISA method (Figure S4, Supporting Information) and achieves a four-log dynamic range **Figure 1d**^33^.

### 2.3. Specificity of the AEM Sensor

The specificity study of the biosensor is important to make sure that the measured signal is indeed from the specific interaction of antibodies with EVs and not from the non-target interactions between antibodies and interfering molecules from the sample^2^. Therefore, several control experiments were performed to address the specificity. Firstly, as control experiments, the anti-CD63 antibody functionalized sensor was incubated with the EV sample and then with silica reporter particles. As shown in **Figure 2a-c**, a large voltage shift was observed from the target sample, despite the low EV concentrations. A large shift was obtained with human plasma with anti-CD63 capture (**Figure 2d**), while plain PBS (**Figure 2e**) and isotype capture (**Figure 2f**) produced relatively low shifts, showing the specificity of our platform obtained from the controlled wash.

**Figure 2.**
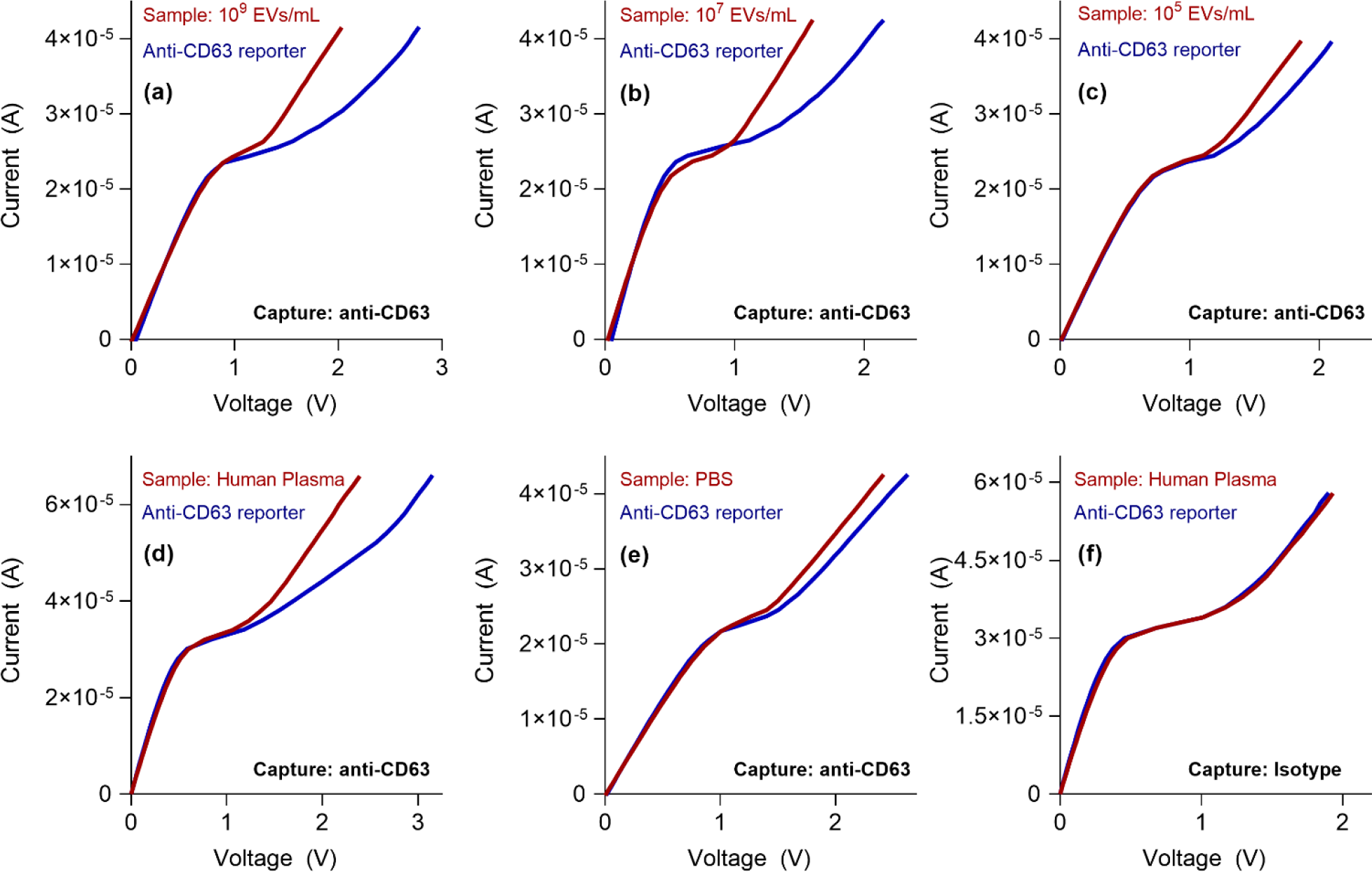
(a), (b), and (c) show the voltage shift with known quantities of EV with anti-CD63 capture and reporter. (d) and (e) use human plasma and PBS respectively as positive and negative controls to show a small shift produced by the EV-free PBS sample. (f) Shift produced by human plasma is negligible with isotype control capture antibody. EVs were isolated from a DiFi cell conditioned medium.

### 2.4. Obtaining Other Subfractions of EVs

Our calibration curves are highly reproducible due to the electrokinetic nature of the signal, eliminating the need for frequent recalibration. A key feature of our platform is the consistent slope observed in the voltage shift versus the logarithm of the (molecular or EV) analyte concentration (slope ∼2*R*_*B*_*T*/*F*), which holds true across various analytes tested, including proteins and lipoproteins. This universality is related to the logarithm dependence of the Zeta potential to surface charge^24, 34^. With our small sensors, we find this calibration curve to be incubation time independent^26, 28^. With small molecules, we do find antibody affinity sensitivity such that the standard curve in the semi-log plot of voltage shift versus concentration translates with different antibody-protein pairs while retaining its slope. However, because of high reporter concentration, irreversible reporter association becomes affinity independent. This would produce a universal calibration curve for all target proteins on the CD63-EVs that are reported by the silica particle. To demonstrate this universality, we use antibodies for tEGFR as a reporter which is known to be present in the vast majority of DiFi-derived EVs showing the voltage response remains the same and coincides with the CD63 calibration plot (**Figure 3a**)^6^.

**Figure 3.**
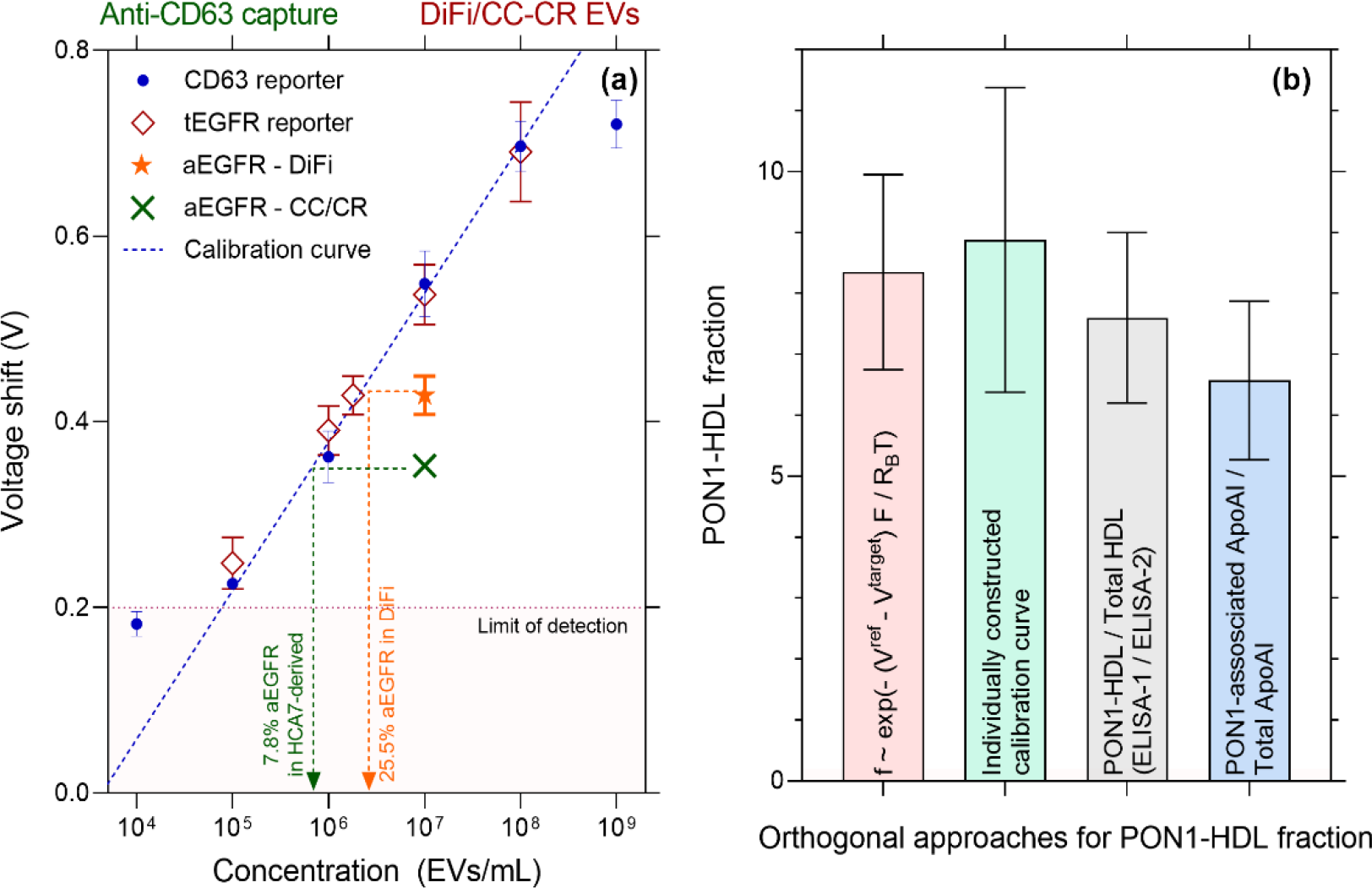
(a) Voltage response with anti-CD63 capture and other reporters. Anti-CD63 reporters and anti-tEGFR reporters give similar responses as they are both present on a vast majority of EVs and because our platform is independent of the affinity of reporter antibody due to the high abundance of reporters. A universal linear calibration between voltage shift and log concentration, corresponding to the linear region of Fig. 1d, is used to estimate the fraction. (b) Four orthogonal approaches using untreated plasma demonstrates that the calibration-free universal correlation for the colocalization fraction *f* = exp (−(*V*^*ref*^ − *V*^*target*^)*F*/2*R*_*B*_*T*), gives a similar estimate as an independently constructed calibration plot (green bar) and two other orthogonal approaches with ELISA-1 and ELISA-2 (grey bar) defined in our previous work^24^. (F is the Faraday constant, T the temperature, and R_B_ the Boltzmann constant from the Boltzmann theory of ion distribution^34^, *V*^*ref*^ and *V*^*target*^ are the voltage shift signal with CD63 and EGFR silica reporters.) Also shown, using a PON1 pulldown and quantifying the ubiquitous protein ApoA1 on the pulled-down volume to normalize against the total ApoA1 (blue bar). HDL was used to verify this universality due to its high concentration and ease of validation using orthogonal methods.

The universal calibration curve of our platform provides a straightforward approach for normalizing EV measurements against a reference species, regardless of the specific subset of EVs of interest. For a given reporter, our earlier studies^24^ indicate that 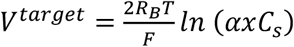 where α is a constant related to the zero potential reference and the valency of the charge, *C*_*s*_ is the concentration of captured species and *x* is the fraction of captured species having colocalized proteins of interest (see definitions in the caption of **Figure 3**). We can similarly write 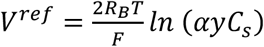 where *y* is the fraction of captured species having reference colocalized protein of interest where the reference could be a ubiquitous protein such as CD63, CD9, or CD81, or even a cocktail. In that case, the percentage of colocalized proteins compared to a reference protein obeys the universal calibration curve *f* = *x*/*y* = exp (−(*V*^*ref*^ −*V*^*target*^)*F*/2*R*_*B*_*T*). It is important that the capture antibody be shared for both reporters (to eliminate *C*_*s*_). For example, CD63 capture and CD63 reporter can be used as a reference for EVs that are captured with CD63 and produces a shift of *V*^*ref*^, while a reporter for a protein of interest produces a signal of *V*^*target*^ (after CD63 capture). This also implies that the lowest percentage one can successfully characterize is the inverse of the dynamic range span. Since we have a four-log dynamic range, we are able to reach 0.01% **(Figure 4b)**.

**Figure 4.**
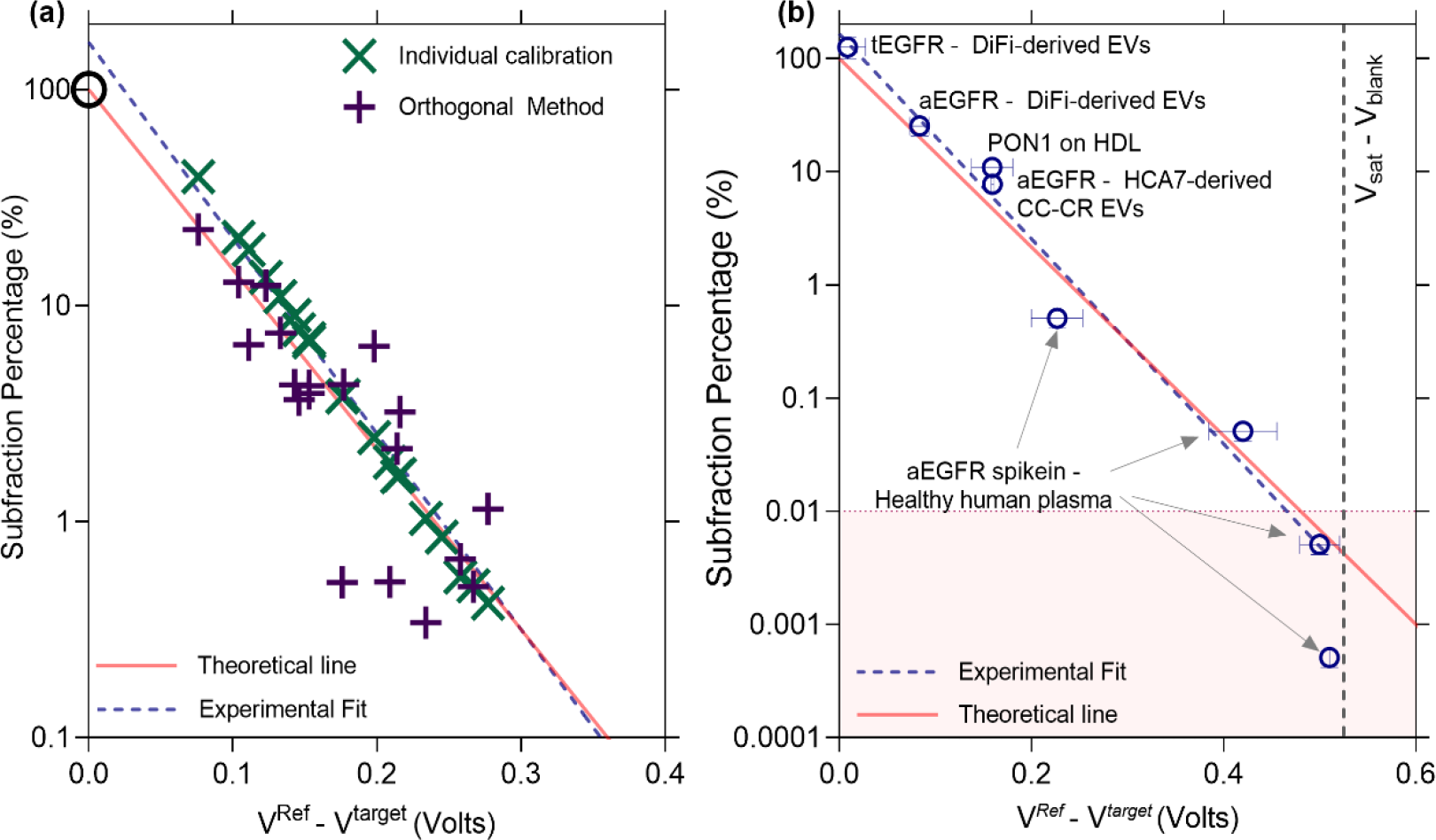
(a) Subfraction percentage estimated from individual calibration curves for PON1-HDL and PON1-free HDL as well as orthogonal method (ELISA-1/ELISA-2) plotted against the difference of *V*^*ref*^ (PON1-free HDL) and *V*^*target*^ (PON1-HDL) using 20 independent human untreated plasma samples. The theoretical line is the universal colocalization fraction *f* = exp (−(*V*^*ref*^ − *V*^*target*^)*F*/*R*_*B*_*T*) where − *F*/*R*_*B*_*T* is the slope. The experimental fit shows a similar slope as that of the theoretical line. (b) aEGFR subfractions determined using anti-CD63 calibration curve (reporter independent as previously shown) and plotted against *V*^*ref*^ − *V*^*target*^ for various capture-reporter antibody combinations and for different cell culture media/plasma. The data again follow the theoretical line to confirm that no calibration is needed. This universality is especially useful when determining non-abundant EV fractions that cannot be calibrated easily due to their unknown concentrations.

To demonstrate the calibration is universal for all CD63-EVs and is independent of the affinity of the capture or reporter antibodies, we use high-density lipoproteins as our model system as they are simpler to characterize using less sensitive orthogonal methods and are in high abundance to use methods such as modified ELISA^26^. In the first approach, using anti-ApoA1 as capture, we obtain two voltage shifts corresponding to the target (anti-PON1 reporter) and reference (anti-ApoA1 reporter). Using the theoretical estimate *f*, we obtain similar values compared to other methods – including constructing individual calibration plots (**Figure 3b)**, yielding that about eight percent of all HDL from healthy donors contains PON1. The two orthogonal methods yield values that are very close to what we obtained using our method (**Figure 3b**). The first orthogonal method, ELISA-1/ELISA-2, is a modified version of ELISA previously used in our work^26^. The second orthogonal method uses the primary protein on HDL (ApoA1) in the immunoisolation and as a reference, with a corresponding voltage shift *V*^*ref*^, and the target PON1 with a voltage shift of *V*^*target*^ to obtain the fraction of ApoA1-HDL that displays PON1. The measured PON1-HDL fraction obeys the same relationship *f* = exp (−(*V*^*ref*^ − *V*^*target*^)*F*/2*R*_*B*_*T*) as EVs. This colocalization calibration should hence be valid for any nanocarrier, EV, HDL, LDL, etc. Additionally, we compare PON1-HDL (target) and PON1-free HDL (reference) for 20 human participants, and we compare the subfraction from individually constructed calibration plots as well as orthogonal methods (ELISA-1 and ELISA-2 in our previous published work) and observe that the same universal curve for EVs also applies to PON1+ HDL (**Figure 4a**) ^26^. We note the larger scatter of the orthogonal methods at low concentrations. These methods are prone to many errors even for simpler lipoproteins since lipid peroxides interfere in the enzymatic reaction (this is described in detail in our previous work^26^).

Using the reporter in high abundance to accommodate low-abundance target proteins, we successfully characterized aEGFR (untethered EGFR and EGFRviii) and determined their relative fractions using CD63 calibration plot as well as the theoretical estimate *f*, as illustrated in **Figure 4b**. We are able to collapse all colocalization data for tEGFR-CD63, aEGFR-CD63 and PON1-ApoA1 from plasma and cell culture media of DiFi and HCA7-derived CC-CR cell lines using the same universal calibration curve. Additionally, to investigate the minimum detectable subfraction colocalization directly in raw plasma, we conducted spike-in experiments using varying quantities of DiFi-media derived extracellular vesicles (EVs) with known aEGFR fractions. These EVs were introduced into healthy pooled human plasma, which initially had minimal aEGFR-EVs content, resulting in the generation of simulated plasma samples with known amounts of aEGFR for our assessment. In **Figure 4b**, we assessed the minimum detectable quantity of aEGFR by analyzing simulated plasma samples containing known amounts of aEGFR fraction and comparing them to the estimated amounts obtained from our platform. The universal calibration curve successfully quantified aEGFR+ CD63 EVs at levels as low as 0.01%. Additionally, various fractions aligned well with our theoretical trend, highlighting that aEGFR fractions can be directly quantified from plasma without the requirement of isolation steps such as ultracentrifugation.

### 2.5. Clinical Sample Analysis

Having established the robustness of the voltage signal for normalized colocalization assay of raw plasma EVs, we test its capability to differentiate patient glioblastoma multiforme (GBM) and healthy plasma samples. The human plasma samples from the twenty GBM cancer patients and five healthy subjects were used for analysis. Their fractions of EVs with aEGFR yield p-value of 3.3E-5 (**Figure 5a**) and 3E-4 (**Figure 5b**) respectively, when normalized with respect to CD63 and tEGFR, with an AUC of 0.99 for both (**Figure 5d**,**e**). The patient samples have an aEGFR-CD63 EV fraction ranging from 0.01 % to 60%, a 4-decade variation. The fraction for healthy samples is close to the LOD of 0.01%, suggesting an undetectable amount of aEGFR by the sensor. In contrast, the fraction of tEGFR EVs in GBM plasma (normalized to CD63 EVs) gave a p-value of 4.97E-2 (**Figure 5c**) and AUC of 0.755 (**Figure 5f**) with a net variation from 25% to slightly over 100% (corresponding to samples that contain EVs with one copy of CD63 and one or more copies of tEGFR). This is a substantial difference from aEGFR measurements that not only spanned several orders of magnitude but was also below the level of detection in healthy patients when normalized relative to CD63 or tEGFR. This result agrees with the previously reported findings that glioblastoma cancer patients express more variant III EGFR EVs^33^, which is a subset of overall active EGFR.

**Figure 5.**
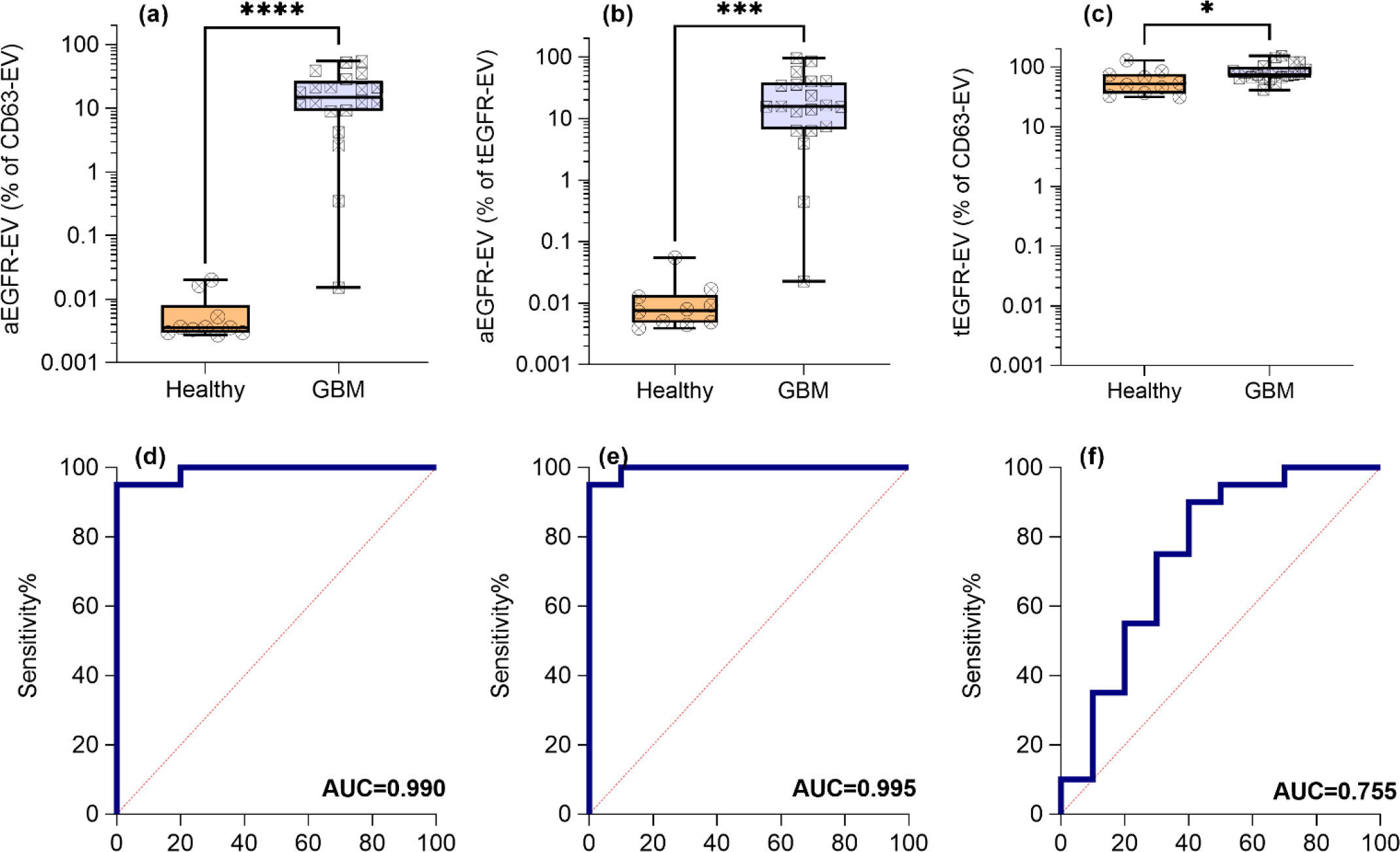
(a,d) The expression and AUC of aEGFR-EVs (normalized to CD63-EVs) with p-value = 3.3E-5, (b,e) aEGFR-EVs (normalized to tEGFR EVs) with p-value = 3E-4 and (c,f) tEGFR-EVs (normalized to CD63-EVs) with p-value = 4.97E-2. All reported p-values were calculated using a parametric test with Welsh’s correction.

## 3. Conclusions

In summary, we have successfully developed an automated, low-cost (<$2 in material), and rapid (< 1 hour) AEM biosensor platform to quantify the fraction of CD63 EVs with a specific cell-surface marker, aEGFR, despite its low abundance and high *K*_*D*_. It is the first colocalization EV assay for untreated plasma. The sensor is optimized for EVs, with proper sensor dimension to reduce incubation time without diminishing sensitivity and robustness. The field from a highly charged silica nanoparticle is amplified by long-range ion depletion action of the ion-selective membrane, to affect the electrokinetics at the membrane surface that controls the ion flux, thus producing orders of magnitude amplification of the voltage signal without distortion despite variations in the EV size. Controlled PBS wash with high drag on the EV and the nanoparticle reporter is used against multivalent adsorption of non-targets to enhance specificity to the extent that high-loss isolation is not necessary and untreated plasma can be used. The AEM microsensor is easy to fabricate and was modified with capture antibody using conventional coupling chemistry. Elimination of analyte loss and irreversible association yielded a normalized calibration curve for the fraction of CD63 EV with a colocalized protein. This universal calibration curve for the colocalization fraction has a linear range between 0.01 to 100 % colocalization (corresponding to 30 to 300000 EVs/μ*L*). EVs quantification of the active EGFR+ EVs in untreated plasma demonstrated that the biosensor is excellent for the detection of specific types of cancer. Glioblastoma clinical sample analysis showed a clear differentiation (AUC∼ 0.99) between cancer patient samples and healthy ones with a single marker colocalization assay, with a p-value of 0.000033 that is superior to any other EV diagnostics.

The current platform could be used to follow possible residual or recurrent GBM and other cancers with amplified aEGFR and tEGFR, as the presence of amplification/mutation of EGFR is frequently retained in metastatic disease. Its use in a screening or diagnostic test to identify cancer type, location, and stage is more complex. Other cancers, like colorectal cancer, can have EGFR amplification and can show an enhanced EGFR signal in plasma from these patients^6^. Therefore, such an EGFR active and total signature might not necessarily indicate the presence of GBM specifically. Likewise, GBM patients can have amplified or mutated EGFR but can also have non-EGFR driven forms of the disease. To establish this as a specific test for a specific cancer would require analyzing a much larger set of GBM cases matched to EGFR status along with testing plasma from other cancers and diseases that might regulate biofluid EGFR. Each distinct tumour type or genotype might regulate the amount of aEGFR, tEGFR, and CD63 EV-carrying forms of such EGFR analytes. Therefore, based on these measures alone it might be possible to tell the difference between different kinds of GBMs and different kinds of diseases that regulate these EV-carried proteins. More broadly, a multi-biomarker version of this platform, which includes markers other than aEGFR or tEGFR and reference EV markers beyond CD63, should enhance the specificity of the screening test. The current diagnostic platform can be scaled up for such large-library testing of untreated plasma from a large cohort of cancer patients to establish specific profiles for different cancers at different stages.

## 4. Experimental Section

### 4.1. Ethics Statement and Human Plasma Samples

We received Glioblastoma and control group samples from Precision for Medicine. An approved IRB protocol is already in place at Precision for the collection of plasma samples from patients. Precision for Medicine works with regulatory authorities and accrediting organizations around the world to ensure that the sample collection process and protocol follow the latest FDA, EMA, and MHRA guidelines. The pooled healthy plasma samples were obtained from Innovative Research (IPLAK2E10ML). Out of 20 GBM plasma clinical samples that we tested, four GBM were purchased from Precision for Medicine and the remaining 16 samples were received from Andrew Scott and Hui Gan, Tumour Targeting Laboratory, ONJCRI, Melbourne, Australia. 10 healthy plasma samples were purchased from Precision for Medicine.

### 4.2. Fabrication of anion-exchange membrane (AEM) sensor

The anion exchange membrane (AEM) composed of polystyrene-divinylbenzene fine particles with strong basic quaternary ammonium groups (R-(CH_3_)_3_N^+^) supported by polyethylene as a binder and polyamide/polyester textile fiber was obtained from the Mega a.s. (Straz pod Ralskem, Czech Republic) and used as a sensor. The AEM sensor was fabricated by embedding a small piece of anion exchange membrane of approximate dimensions 0.3 mm X 0.9 mm in an epoxy resin (TAP Quik-Cast, Tap plastic) using an optimized reported protocol^28^. Briefly, the membrane was hand-cut and placed on the tip of a silicone mold. A glass slide was then placed on top of the membrane piece. A two-component (side A and side B, 1:1 ratio) epoxy polyurethane resin mixture was then pushed to embed the AEM sensor^28^. A 3D-printed sensor reservoir was then attached to the sensor disc using the same polyurethane resin mixture. The prepared sensor was soaked in de-ionized (DI) water overnight before further use.

### 4.3. Fabrication of biochip

A 25 mm × 54 mm (w × l) size biochip having a microfluidic channel of 3 mm × 35 mm × 250μm (w × l × h) was fabricated as shown in **Figure 1**. Three layers of polycarbonate sheet of 0.3 mm thickness were used to fabricate using our reported protocol^28^. Briefly, the sheets with orifices for the inlet, outlet, sensor, and channel were cut using a cutting plotter (FC700. Graphtec Corp., Japan). The sheets were then thermally bonded in an oven (Fisher Scientific, Isotemp Oven) at 177 °C for 15 min. The small pieces of tubes for the inlet and outlet and three different-sized tubes were attached in between the inlet and outlet for mounting a sensor and different electrodes using Acrifix UV glue. The electrode reservoirs were filled with 2% agarose gel to create a barrier between the microfluidic channel and the reservoirs.

### 4.4. Functionalization of the antibody on the AEM surface

The surface modification of the anion-exchange membrane and antibody functionalization followed a previously optimized protocol^28^. Initially, the membrane surface was treated with a 0.1M solution of 3,3’,4,4’-Benzophenonetetracarboxylic acid (Sigma-Aldrich, USA) at pH 7 for 10 minutes. Subsequently, the surface was exposed to UV light at 365 nm (using an Intelli ray 600 shuttered UV floodlight) for 90 seconds while purging with N2 gas. The sensor was then rinsed with deionized (DI) water. This surface modification process was repeated three times to ensure the generation of an adequate number of –COOH groups on the membrane surface. Following that, the sensors were immersed in DI water at pH 2 for 5 hours and subsequently washed with 0.1X PBS at pH 7. The carboxylated membrane surface was then ready for functionalization with the antibody probe using EDC (Thermo Fisher, USA) coupling chemistry. A 20 μL solution of 0.4 M EDC in 50 mM MES buffer at pH 6 was applied to the sensor surface for 40 minutes. The solution was then removed, and the sensor was washed with 1X PBS. Finally, a solution of the probe antibody (0.1 mg/mL) was incubated overnight on the sensor surface at four-degree celcius^28^.

### 4.5. Conjugation of antibody to silica reporter particles

To prepare the carboxylated silica particles, 500 μl of 2.5% silica particles with a size of 50 nm (Microspheres-Nanospheres (New York, USA)) were mixed with 500 μl of 1X PBS. The mixture was then centrifuged at 17000 g for 10 minutes, and the supernatant was discarded. This washing process was repeated twice. After the final wash, the particles were resuspended in 1 ml of 50 mM MES buffer at pH 6. Next, a solution containing equal volumes of 200 mM EDC and 200 mM sulfo-NHS (EMD Millipore, USA) prepared in MES buffer was added to the silica particles. The mixture was stirred for 1 hour at room temperature. Subsequently, the silica particles were washed three times with 1X PBS by centrifugation at 17000 g for 10 minutes each time. Finally, 20 μL of a 0.1 mg/mL solution of the detection antibody was added to the 1 mL of silica particles in 1X PBS. The mixture was mixed overnight at 150 RPM at 4 °C. Afterward, the particles were washed three times with 1X PBS to remove any unbound antibodies from the solution.

### 4.6. Labelling of anti-CD63 Antibody

To confirm the binding of antibodies on the sensor and silica reporters, the anti-CD63 antibody was labelled using Zip Alexa Fluor 488 rapid antibody labelling kit (ThermoFisher, USA) according to the manufacturer’s instructions with some modifications. In brief, 10 μl of 1M sodium bicarbonate solution was added to the 100 μl of 0.5 mg/mL antibody solution. The antibody solution was then added to the Alexa Fluor dye. The solution was mixed using a micropipette and incubated for 15 min at room temperature. The free dye was removed using Amicon Ultra 100k centrifugal filter devices and the labelled antibody was stored at 4 °C till further used.

### 4.7. Voltage signal measurements

The current-voltage characteristics (CVC) signal of the AEM sensor was evaluated using a Gamry Potentiostat/Galvanostat/ZRA (Reference 600, Gamry Instruments Inc., USA) in a four-electrode configuration. Crocodile clips were used to connect the electrodes to the instrument, and they were securely mounted on the biochip. The current was supplied through the two platinum electrodes, while the potential across the membrane was measured using Ag/AgCl reference electrodes (World Precision Instruments, USA). CVC measurements were conducted in a 0.1X PBS solution. The current was incrementally increased from 0 μA to double the limiting current, and the potential was recorded at a step rate of 1 μA/s. The measurements were performed using Gamry Framework software, and the obtained spectra were analysed using Gamry Echem Analyst software. To introduce different buffers into the biochip, a custom-made microfluidic pump was utilized. The pump, controlled by a computer, allowed for the selection of various buffers and adjustable flow rates. A visual representation of the pump is presented in **Figure 1**.

### 4.8. Isolation of Extracellular Vesicles from Cell Culture Media

To optimize different sensing parameters for EV detection, EVs were isolated from the human colorectal cancer cell line, DiFi using ultracentrifugation. Human DiFi cells were cultured as previously described^32^. The cell culture media was centrifuged at 250g and then 2500g both for 10 min in order to remove cellular debris and the supernatant was collected in a new vial. Then the supernatant was filtered through a 0.22μm polyethersulfone filter (Nalgene) to remove the microparticles by gravity flow. Further, the collected filtrate was concentrated by a 100,000 molecular-weight cutoff (Millipore) centrifugal concentrator. Finally, the high-speed centrifugation at 167,000g for 4 hours in an SW32 Ti swinging-bucket rotor (Beckman Coulter) was used to separate and further wash the EV from the concentrate collected in the previous step. The EVs suspended in PBS 25mM HEPES were used immediately or stored at -80 °C in aliquots for further use. The average particle size and particle concentration were measured using the nanoparticle tracking analysis (NTA) method (NanoSight™ NS300, Malvern Instruments Ltd., UK). Similar approach was taken for HCA7-derived CC-CR cell line (colorectal cancer cell line).

### 4.9. Antibodies

Purified mouse anti-human CD63 antibody (Catalog No. 556019) and isotype control antibody (Catalog No. 555746) were bought from BD Biosciences, USA. Mab806 antibody (ABT-806, Catalog No. TAB-228CL) and Human EGFR (cetuximab) antibody (Catalog No. MAB9577) were purchased from Creative Biolabs, USA, and R&D Systems respectively.

### 4.10. ELISA

Maxisorp 96-well plate (Nunc) was coated with an anti-CD63 antibody by incubating overnight at 4 °C. Then, the plate was washed with 1X PBS. The plate was blocked using 2% BSA in a blocking solution for 1 hour at room temperature. EV samples of 100 μL were added to each well and incubated for 2.5 hours at room temperature. After washing, 100 μL of biotinylated anti-CD63 antibody was added to each well for 1 hour at room temperature. Then, 100 μL of HRP-Streptavidin molecules were added to each well and incubated for 45 min at room temperature. The absorbance signal was measured at 450 nm using a plate reader.

## Supporting Information

Supporting information is available from the Wiley Online Library or from the authors.

## Acknowledgments

This work was partially supported by the NIH Commons Fund, through the Office of Strategic Coordination/Office of NIH Director, 1UH3CA241684-01 (HCC & SS) and 1UH3CA241685 (RJC).

## Data Availability Statement

The data that support the findings of this study are available from the corresponding authors upon reasonable request.

