## Supplementary Material for "Detection of EGFR and its Activity State in Plasma CD63-EVs from Glioblastoma Patients: Rapid Profiling using an Anion Exchange Membrane Sensor"

---

The supplemental information provides additional details on the following.

**Figure S1.** Photograph of (a) cut anion-exchange membrane (AEM), (b) top and bottom parts, and (c) fully prepared AEM sensor.

**Figure S2**. (a) AEM sensor functionalization using Alexa fluor 488 labeled CD63 antibody and (b) Silica reporter particles functionalization using Alexa fluor 488 labeled CD63.

**Figure S3**. NTA spectrum of isolated EVs. The average hydrodynamic diameter and concentration of the EVs were found to be 117 nm and $1 \times 10^{10}$ particles/mL, respectively.

**Figure S4.** CD63 ELISA of different concentrations of isolated DiFi EVs.

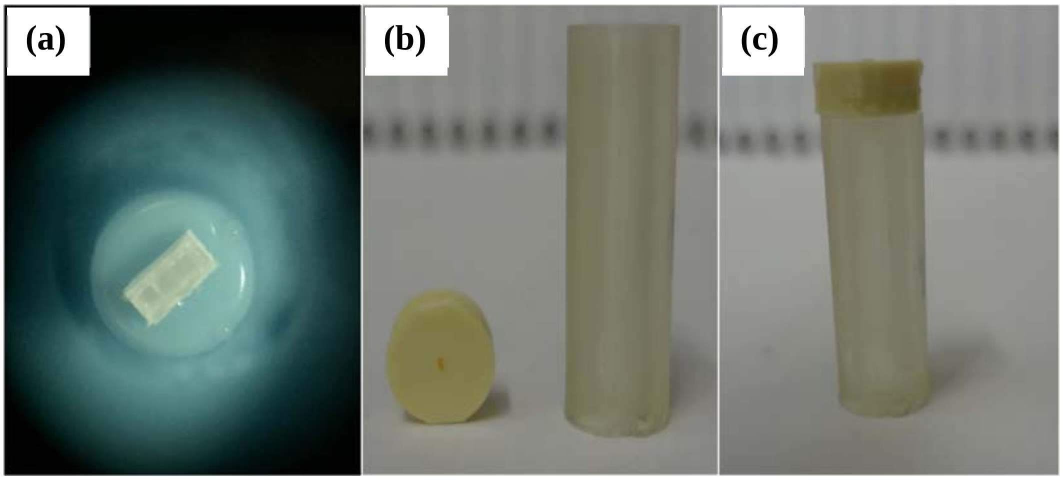


**Figure S1.** Photograph of (a) cut anion-exchange membrane (AEM), (b) top and bottom parts, and (c) fully prepared AEM sensor.

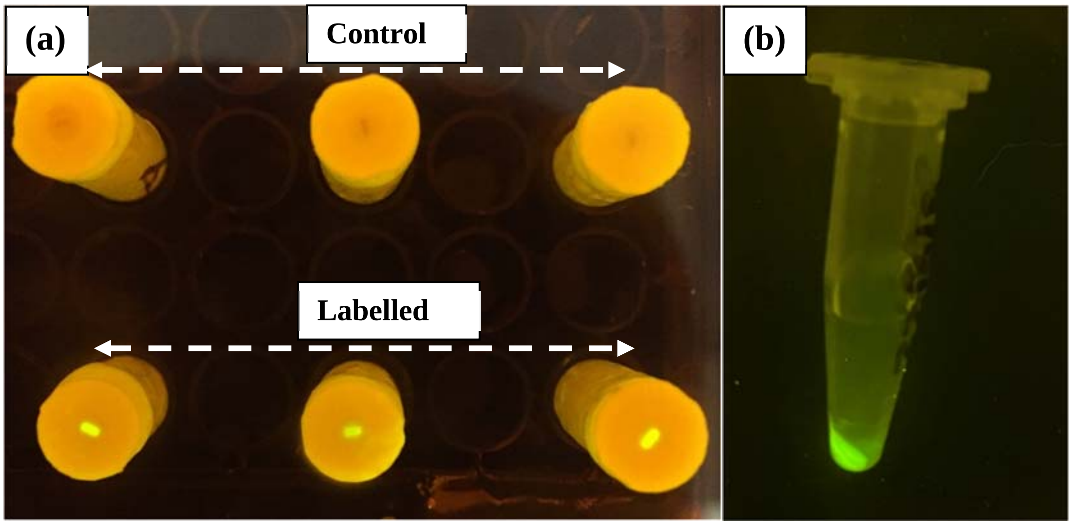


**Figure S2**. (a) AEM sensor functionalization using Alexa fluor 488 labeled CD63 antibody and (b) Silica reporter particles functionalization using Alexa fluor 488 labeled CD63.

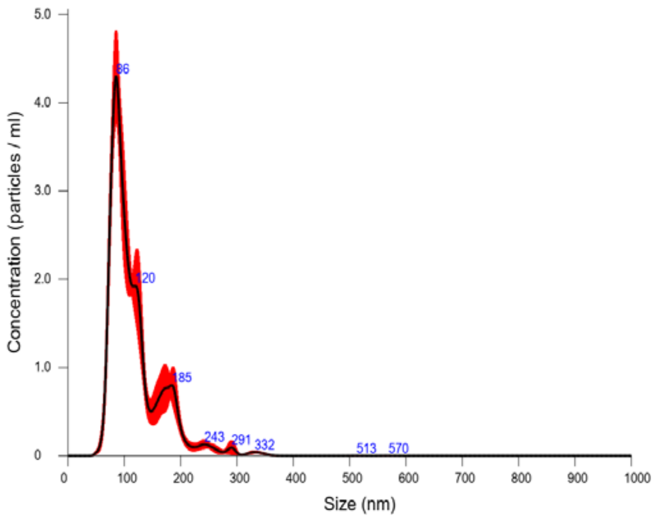


**Figure S3**. NTA spectrum of isolated EVs. The average hydrodynamic diameter and concentration of the EVs were found to be 117 nm and $1 \times 10^{10}$ particles/mL, respectively.

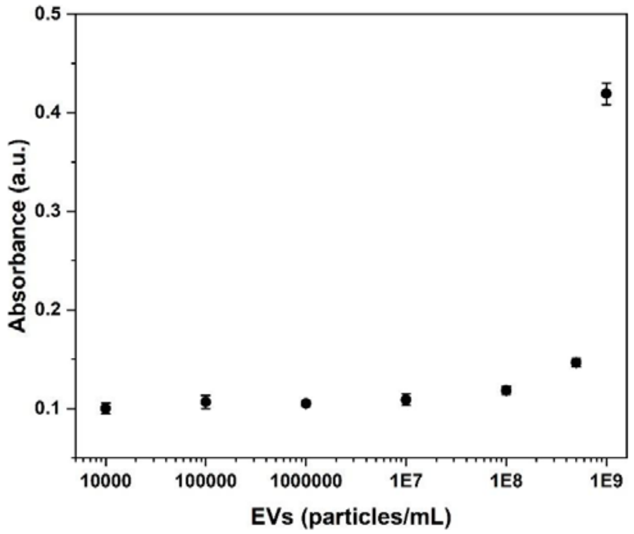


**Figure S4.** CD63 ELISA of different concentrations of isolated DiFi EVs.
